# Mechanistic dissection of alga recognition and uptake in coral-algal endosymbiosis

**DOI:** 10.1101/2022.12.13.520278

**Authors:** Minjie Hu, Yun Bai, Xiaobin Zheng, Yixian Zheng

## Abstract

Many corals form a mutually beneficial relationship with the dinoflagellate algae called *Symbiodiniaceae*. Cells in the coral gastrodermis recognize, phagocytose, and house the algae in an organelle called symbiosome, which supports algae photosynthesis and nutrient exchange with corals^1–3^. Rising ocean temperature disrupts this endosymbiotic relationship, leading to alga loss, coral bleaching and death, and the degradation of marine ecosystems^4–6^. Mitigation of coral death requires a mechanistic understanding of coral-algal endosymbiosis. We have developed genomic resources to enable the use of a soft coral *Xenia species* as a model to study coral-algal endosymbiosis^7^. Here we report an effective RNA interference (RNAi) method and its application in the functional studies of genes involved in early steps of endosymbiosis. We show that an endosymbiotic cell marker called LePin (for its <u>Le</u>ctin and kazal <u>P</u>rotease <u>in</u>hibitor domains) is a secreted lectin that binds to algae to initiate the formation of alga-containing endosymbiotic cells. The evolutionary conservation of LePin among marine endosymbiotic anthozoans suggests a general role in coral-algal recognition. Coupling bioinformatics analyses with RNAi and single cell (sc)-RNA-seq, we uncover three gene expression programs (GEP) influenced by LePin during the early and middle stages of endosymbiotic lineage development. Further studies of genes in these GEPs lead to the identification of two scavenger receptors that support the formation of alga-containing endosymbiotic cells, most likely by initiating phagocytosis and modulating coral immune response. We also identify two actin regulators for endosymbiosis, which shed light on the phagocytic machinery and a possible mechanism for symbiosome formation. Our findings should usher in an era of mechanistic studies of coral-algal endosymbiosis.

## Main

The importance of endosymbiosis for corals has been recognized for many decades^8,9^, but the biology of this mutually beneficial relationship remains obscure because it has been subject to limited molecular studies. The alarming loss of coral reefs and the associated degradation of marine environments have galvanized efforts to understand coral-algal endosymbiosis in recent years. Multiple bulk RNA-seq comparisons between apo-endosymbiotic and endosymbiotic corals or control and bleached corals have identified a large number of differentially expressed genes. Although these genes could be candidate regulators for endosymbiosis, many of them, such as the NFkB pathway genes and lectin genes^10–13^, may be expressed in different cell types and have general functions beyond endosymbiosis. Their differential expression may represent a secondary response to alga loss. Through developing *Xenia* as a model organism for studying coral endosymbiosis, we have identified different cell types based on scRNA-seq and established methods to characterize lineage progression of the endosymbiotic cell type^7^. Among the genes highly expressed in *Xenia* endosymbiotic cells, 89 are markers for the endosymbiotic cell lineage because they are specifically expressed in this cell type, while an additional 686 highly expressed genes in endosymbiotic cells are also found in a few other cell types^7^.Interestingly, some of these genes are conserved and specifically expressed in the endosymbiotic cells of a stony coral as revealed by a recent study^14^. Functional analyses of these genes could yield mechanistic insights of coral endosymbiosis. Such insights are important for understanding how corals bleach and it may aid in developing mitigation strategies. For example, recent studies suggest that corals harboring heat resistant algae are more resistant to bleaching than those corals selecting the non-heat resist endosymbionts^15–18^. Identifying the algal selection mechanism could help engineering heat resistant corals.

### RNA interference as an efficient means for gene down regulation in *Xenia sp*

Our genomic study has found key components of RNA interference (RNAi) machinery in *Xenia*^7^. Since corals are known to actively take up fluids from the environment through macropinocytosis^19^, simple incubation of *Xenia* with RNAi oligos could result in uptake of the oligos and RNAi-mediated gene silencing. We tested this by incubating *Xenia* polyps with media containing Cy3-labeled oligo-T for a day and found strong Cy3 signals in *Xenia* tissues (Fig. 1a). We next synthesized control short hairpin RNA (shRNA) and shRNAs targeting *Xenia* genes predicted to encode tubulin, GAPDH, and lamin-B2 (Supplementary Table S1), and incubated them with *Xenia* polyps. As shown by Real-Time Quantitative Reverse Transcription PCR (RT-qPCR), the shRNAs substantially reduced the expression of these genes compared to the control (Fig. 1b). Thus, the RNAi approach should allow us to establish the function and dissect the mechanism of candidate genes required for endosymbiosis.

**Fig. 1.**
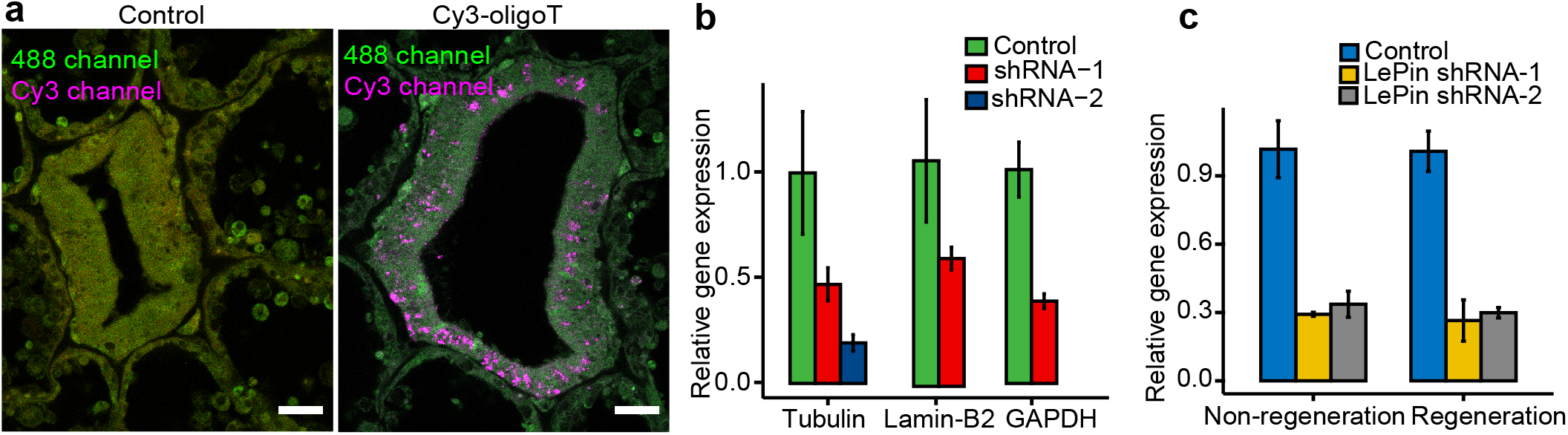
shRNA knockdown of genes in *Xenia*. **a**, Incubation of *Xenia* polyps in artificial sea water containing Cy3-labeled oligoT (20 nucleotides in length, displayed in magenta) resulted in an uptake of the labeled oligos into tissues. The background signal in the 488 channel (displayed in green) was used to help visualize the outline of tissue sections. Scale bar, 20μm. **b,c**, shRNA-mediated knockdown of genes predicted to encode for tubulin, lamin-B2, GAPDH (**b**), and LePin (**c**). For LePin, shRNA is effective for both unperturbed polyps and tentacle amputated polyps undergoing regeneration. Gene expressions are quantified by qRT-PCR. The number of shRNA used for each gene is indicated. Three different polyps were used for each shRNA treatment.

### LePin aids the formation of new alga-containing endosymbiotic cells

During lineage progression *of Xenia* endosymbiotic cells, the formation of alga-containing cells begins when progenitor cells select and bind to the correct *Symbiodiniaceae* species. Lectins are a family of pattern recognition proteins known to exhibit binding specificity to different glycans on microbial surfaces^20^. They have been implicated in alga selection/recognition during coral endosymbiosis^21–23^. Finding the lectin specifically involved in selecting/recognizing the preferred endosymbiotic algae among numerous lectins in a given cnidarian genome^24^ has been challenging. We previously identified expression of a lectin called LePin as a marker for the *Xenia* endosymbiotic cell type^7^. Using sequence analyses, we found LePin-like proteins with similar domain compositions in the endosymbiotic marine anthozoans, while the most similar lectins in a non-endosymbiotic marine anthozoan, *Nematostella*, lacked the C-lectin and EGF domains (Extended Data Fig. 1). This suggests that LePin could be the lectin involved in initiating endosymbiosis by recognizing proper endosymbiotic algae for corals. To study the role of LePin, we designed two shRNAs targeting different regions of LePin (Extended Data Fig. 2a). *LePin* expression was efficiently reduced by both shRNAs in intact *Xenia* polyps or polyps undergoing regeneration after tentacle amputation^7^ (Fig. 1c). Since *LePin* is strongly and specifically expressed in the endosymbiotic lineage in *Xenia*, its down regulation is specific in cells of this lineage.

Next, we performed scRNA-seq using the regenerating *Xenia* polyps treated by control or LePin shRNA (Extended Data Fig. 3). Previously, we established the stages of endosymbiotic lineage progression using trajectory analyses based on scRNA-seq of either full grown *Xenia* polyps or polyps undergoing regeneration after tentacle amputation^7,25^. Using scRNA-seq, we established gene expression patterns of the endosymbiotic cells as they progress from alga-free progenitors to alga-carrying cells, and finally to cells with gene expression patterns suggesting that they are post endosymbiotic (algae free)^7^. This lineage progression was supported by ethynyl deoxyuridine (EdU) pulse-chase to track the newly divided *Xenia* cells that take up algae^7^. We constructed the trajectory of endosymbiotic cell lineage progression based on the scRNA-seq of the shRNA-treated samples and analyzed the cell distribution along the trajectory (Fig. 2a). We then utilized our previously identified genes enriched in the progenitor endosymbiotic cells^7^ and mapped them in the new trajectory. Most of the genes showed highest expression in the progenitor cells in both control and *LePin* RNAi-treated samples (Fig. 2b, Extended Data Fig. 4), confirming the direction of lineage progression (Fig. 2a). The major peaks in the control and *LePin* shRNA-treated samples represent the alga-containing endosymbiotic cells because their gene expression patterns are highly similar to the transcriptome of the *Xenia* alga-containing cells isolated by Fluorescence Activated Cell Sorting (FACS)^7^. *LePin* RNAi resulted in an increased accumulation of cells prior to the major peak (Fig. 2a, yellow arrow). We then applied the EdU pulse-chase to trace the newly formed endosymbiotic cells in the regenerating *Xenia* treated by control or *LePin* shRNA (Fig. 2c). Using our previously established FACS analysis method^7^, we found a significant reduction in the newly divided cells (EdU+) that contained algae upon *LePin* RNAi (Fig. 2d). Thus, LePin supports the formation of new alga-containing endosymbiotic cells.

**Fig. 2.**
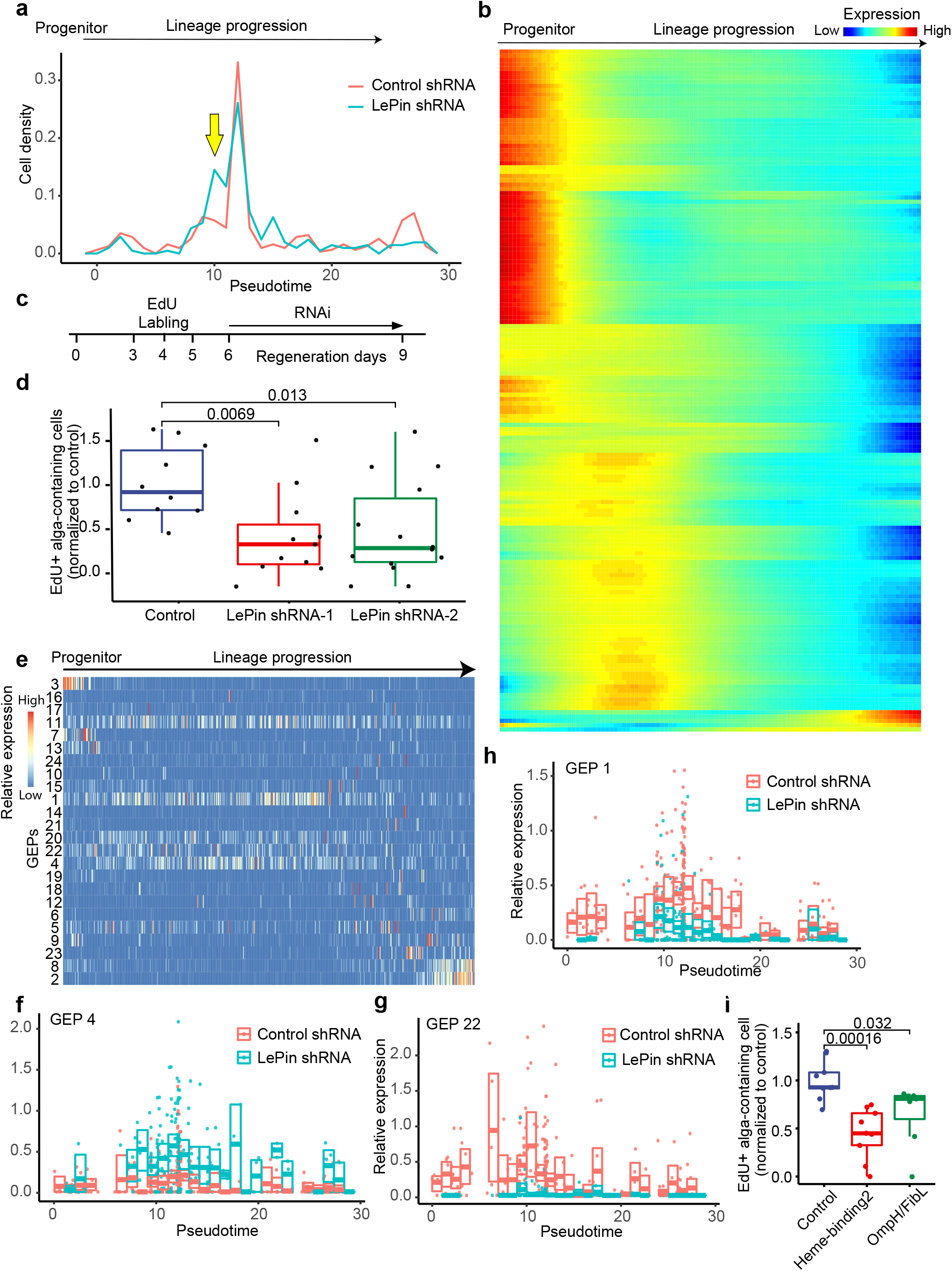
LePin aids the formation of new alga-containing cells and impacts gene expression in the endosymbiotic cell lineage. **a**, Endosymbiotic cell distribution along the lineage developmental trajectories based on the scRNA-seq of control and LePin shRNA-treated regenerating polyps. The pseudotime is calculated by Monocle and used to display the lineage trajectory. Yellow arrow indicates a population of cells accumulated prior to the major peak upon LePin RNAi. **b**, Heat map of gene expression along the developmental progression as measured by scRNA-seq. Most previously defined genes expressed in the progenitor endosymbiotic cells (not carrying algae) have higher expression during the early stages of lineage progress than later stages in the new trajectory analysis. **c**, Experimental design for tracking the newly formed alga-containing endosymbiotic cells. EdU was used to label newly divided cells during day 3 and 4 of *Xenia* regeneration. After EdU washout, shRNA was added on day 6 followed by FACS based quantification of EdU positive (+) and alga-containing cells. **d**, Quantification of the number of newly formed (EdU+) alga-containing endosymbiotic cells upon LePin RNAi. Each animal is normalized against the mean of the control shRNA within the same batch of experiment. Four batches of independent experiments were performed. Each dot stands for an animal. A total of 10-13 regenerating polyps were used for each shRNA treatment. **e**, The GEPs identified by NMF analysis. The cells are ordered along the trajectory of lineage progression. The relative expression of GEPs in each cell is indicated by the heatmap. **f-h**, The expression of GEPs 1 (**f**), 4 (**g**), 22 (**h**) in each cell along the lineage trajectory. Each dot stands for a cell in control or LePin shRNA treated samples. The box plot indicates the GEP expression (see Method) distribution for the cells within one pseudotime unit. Lines in the box represent the median; the upper and lower edges of the box represent the upper and lower quartiles, respectively. **i**. Knocking down Omph/FibL and Heme-binding2 in GEP 1 leads to the reduction of the newly formed alga-containing cells. Quantification as in (**d**). Each dot stands for an animal. 7-9 animals were used for each treatment. The p values of the unpaired Student’s t-test are indicated in **d** and **i**.

### LePin influences three gene expression programs containing candidate genes for early and mid-stages of endosymbiotic lineage progression

As a lectin, LePin could be involved in initiating alga selection, recognition, and subsequent steps of endosymbiosis. If it served any of these roles, elimination of LePin would be expected to impact gene expression programs of the endosymbiotic cells. Applying non-negative matrix factorization (NMF) to scRNA-seq data, it is possible to computationally identify co-expressed genes in individual cells, called metagenes or gene expression programs (GEPs)^26–28^. GEPs may govern cell type identities and/or cell activities^28^. Therefore identifying the GEPs with altered expression upon LePin knockdown could facilitate the discovery of genes involved in endosymbiosis. We applied NMF to the endosymbiotic cell lineage and identified 24 GEPs (Supplementary Table S2). We then coupled the lineage trajectory with the NMF analysis to uncover the GEP expression levels of the 24 GEPs in individual cells during endosymbiotic lineage development (Fig. 2e). We reasoned that comparisons of the expression of GEPs in control and *LePin* RNAi-treated samples along the lineage progression may allow us to identify the candidate GEPs that regulate endosymbiosis (Supplementary Table S3). Among the GEPs that showed expression changes upon *LePin* RNAi, we found three, GEP1, 4, and 22, that showed changes during the early and middle stages of endosymbiotic cell lineage development (Fig. 2f-h). This suggests that the three GEPs contain genes that function at a relatively early stage of endosymbiosis and are affected by LePin function. Since *LePin* is one of the top 30 weighted genes that contribute most to the GEP1 (Supplementary Table S2), we randomly selected two genes in this GEP based on the gene IDs, Xe_016304 and Xe_011936. Xe_016304 is predicted to encode a Heme binding protein (Heme binding protein 2_1), while Xe_011936 is predicted to encode a protein containing two domains, OmpH (Outer membrane protein H) and FReD (Fibrinogen-related domain, FibL) and we referred to this protein as Omph/FibL^7^ (Extended Data Fig. 2b). We previously identified the *Omph/FibL* gene as one of the 89 endosymbiotic cell markers, while Heme binding protein 2_1 (*Heme-binding2*) is one of the 686 highly expressed genes in the endosymbiotic cells^7^. RNAi of each of the two genes resulted in a significant reduction of newly formed alga-containing endosymbiotic cells (Fig. 2i). Thus, the three GEPs contain promising candidates to be involved in mediating early steps of endosymbiosis.

### LePin binds to algae and mediates their interaction with *Xenia* gastrodermis

To further understand the role of LePin in alga recognition, we analyzed *Xenia* tissue sections cut across tentacles, the region connecting tentacles to the mouth (tentacle_mouth), the region connecting the stalk to the mouth (stalk_mouth), and stalks (see the illustrations in the inserts in Fig. 3a). As indicated by nuclear staining, the mouth tissue has an opening in the center surrounded by densely packed cells compared to all the other regions in the polyp (Fig. 3a, M, mouth outlined by white dotted lines). We found numerous algae at or near the gastrodermis surfaces of these regions (Fig. 3a). We next developed an unbiased and high throughput method (Extended Data Fig. 5 and Method) to quantify the number of algae at the surface of gastrodermis throughout the tissue sections from the regenerating *Xenia* treated by control or *LePin* shRNA. We found that the *Xenia* stalk_mouth region showed a significant reduction of algae attachment upon *LePin* RNAi (Fig. 3b). Thus, LePin may initiate algae uptake by supporting the interaction between algae and the gastrodermis cells in the stalk-mouth region.

**Fig. 3.**
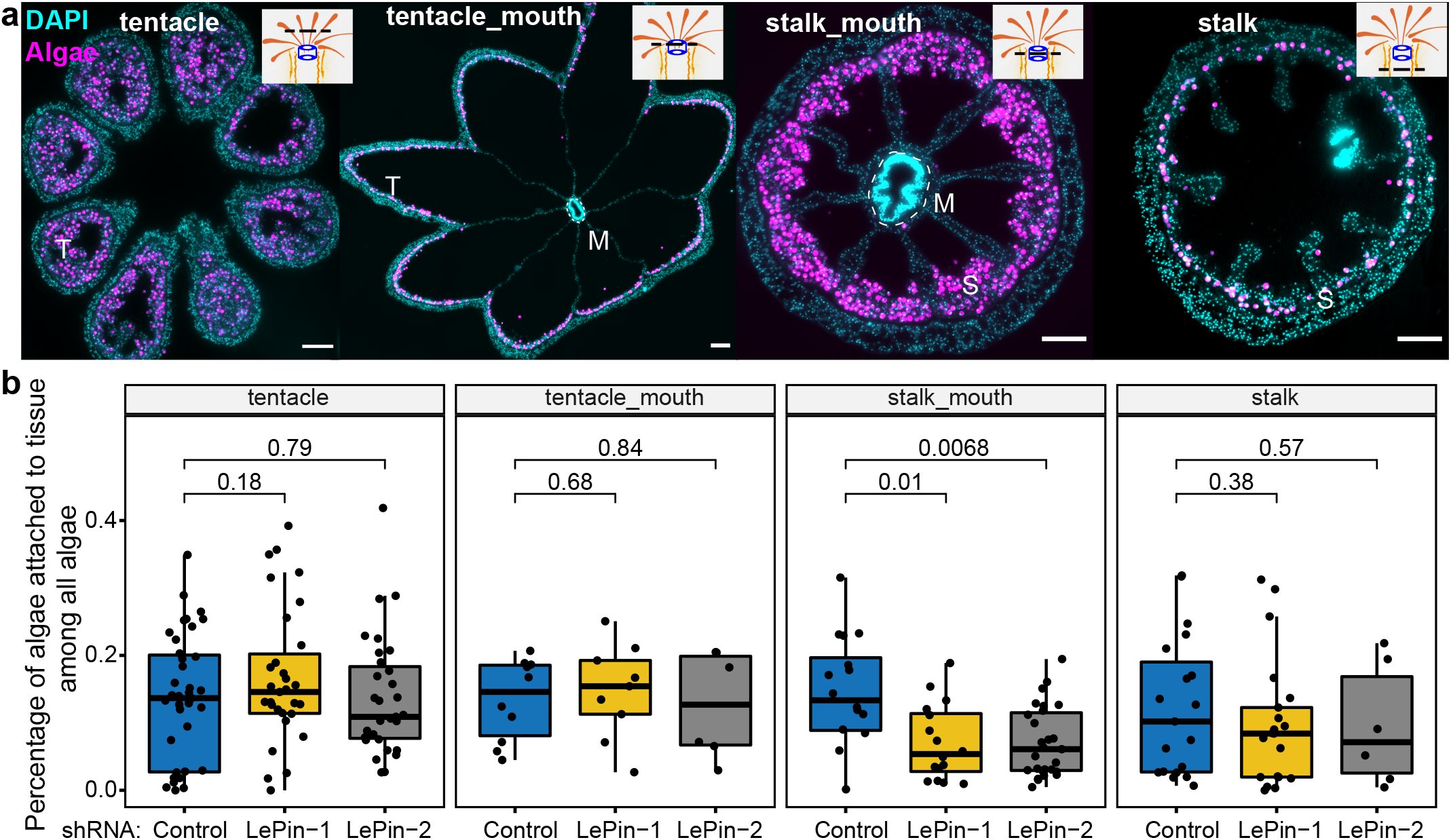
LePin facilitates alga attachment to the stalk_mouth region of the gastrodermis. **a**, Single-plane epifluorescence images illustrating different tissue section categories. Algal cells were imaged by their autofluorescence and are displayed in magenta. Nuclei were stained with DAPI and are displayed in cyan. The black dashed lines in the polyp illustrations at the upper right corners of each panel indicate the positions of individual tissue sections. T, tentacle, M, mouth (outlined by white dotted circles), S, stalk. Scale bar, 100μm. **b**, Reduction of the number of algae attached to the tissues in the stalk_mouth region upon LePin RNAi. 12 regenerating polyps were used per shRNA treatment. Each dot stands for a tissue section. The p values of the unpaired Student’s t-test are indicated.

To further understand how LePin mediates alga recognition, we analyzed the LePin sequence and uncovered a signal peptide at its N-terminus^29^, but we found no transmembrane domains^30^ (Extended Data Fig. 6). This suggests that LePin is a secreted lectin. We next produced antibodies to peptides corresponding to two regions of LePin (Extended Data Fig. 2) and affinity purified the antibodies using the peptides. We performed LePin antibody staining of *Xenia* tissue sections and found LePin on the surface of the mouth tissues (Fig. 4a). RNAi of LePin resulted in a significant reduction of these LePin signals (Fig. 4a top right panels and 4b), demonstrating that LePin is present on the surface of mouth tissue. This is consistent with LePin being a secreted lectin. Since the mouth region of a scleractinian coral, *fungia scutaria*, has been implicated in mediating endosymbiosis establishment in the larvae^31^, the LePin on the *Xenia* mouth surface could aid the recognition/selection of algae as they enter into the gastrodermis.

**Fig. 4.**
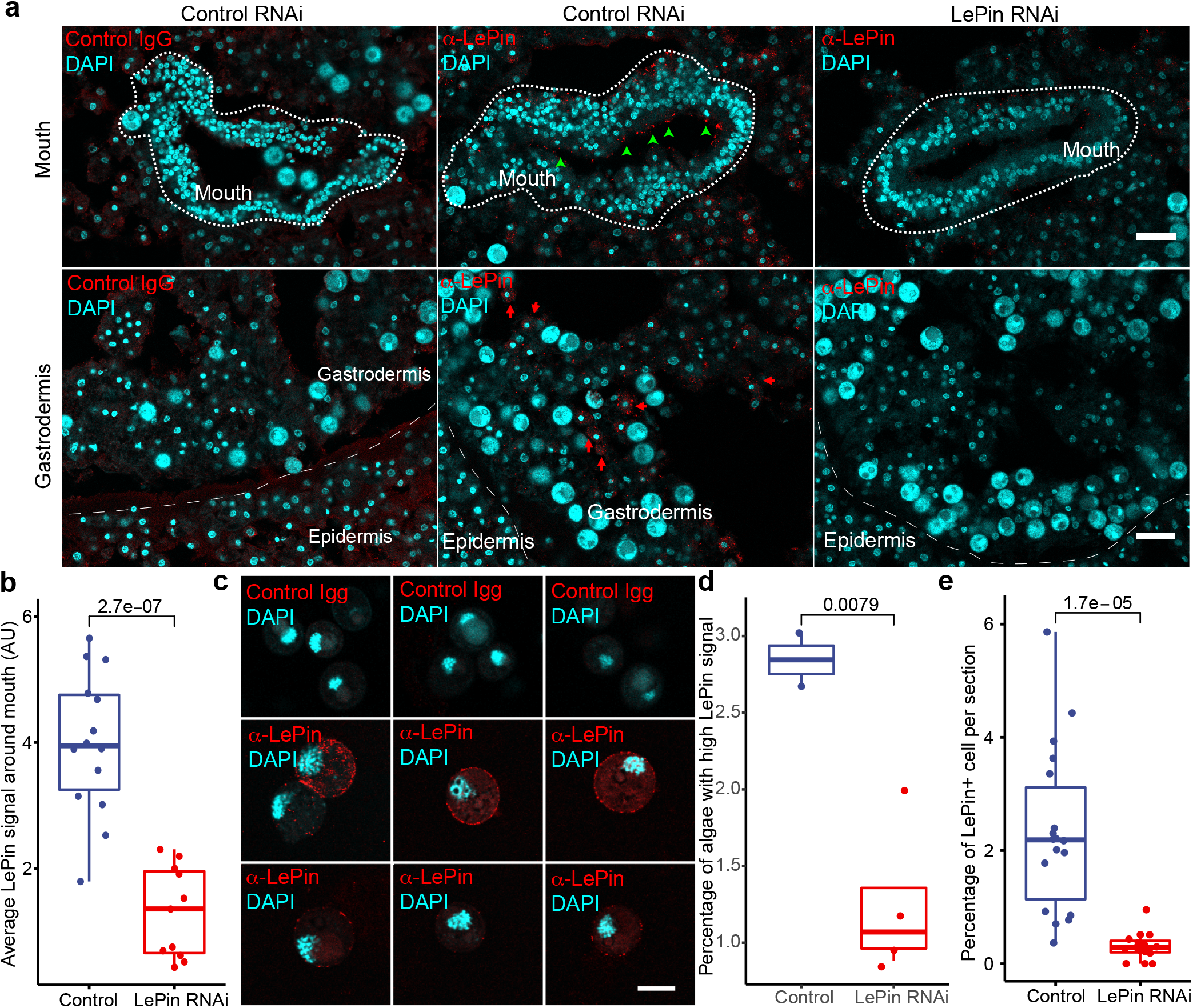
LePin in alga binding and recognition. **a,** Single-plane epifluorescence images illustrating control rabbit IgG or LePin antibody staining in the mouth (top) and gastrodermis (bottom) within the stalk_mouth region. Nuclei are stained with DAPI and displayed in cyan, while the LePin immunofluorescence signal is displayed in red. Green arrows indicate LePin signals around the mouth tissue. Red arrows indicate LePin signal in alga-free cells. Scale bar, 20μm. **b**, Quantification of LePin fluorescence signal intensity around the mouth structure. **c**, Control or LePin antibody staining of the free algae isolated from *Xenia*. LePin showed a range of immunostaining intensity. Scale bar, 10μm. d, Quantification of the percentage of isolated algae with high LePin signal by FACS (see Extended data Fig. 7 for detailed FACS gating and quantification). Each dot stands for a polyp. **e**, Quantification of the percentage of LePin positive and alga negative cells in all the cells in the tissue sections of the stalk_mouth region. The p values of the unpaired Student’s t-test are indicated in **b, d, e**.

To understand how LePin could help alga recognition/selection, we treated the regenerating *Xenia* polyps with control or *LePin* shRNA and then isolated the free algae from these animals. Immunostaining using control and LePin antibodies revealed various levels of LePin signals on the surface of these algae (Fig. 4c). We performed FACS-based unbiased quantification of the LePin signal on algae. Since the algae autofluorescence precluded detection of weak LePin signals on algae, we focused on algae with strong LePin signals. We found that *LePin* shRNA significantly reduced the percentage of algae with strong LePin signals (Fig. 4d, Extended Data Fig. 7). Therefore the secreted LePin can recognize and bind free algae.

Since LePin RNAi caused a significant reduction of algae attached to the gastrodermis of the stalk-mouth region, we next focused on the analysis of LePin in this region. Although a large number of algae in this region made it difficult to detect LePin signal due to a high background signal of algal autofluorescence, we were able to detect clear LePin signals in some cells that do not appear to contain algae and the signal was significantly diminished by *LePin* shRNA (Fig. 4a, bottom panels and 4e). Together, these findings support a role of LePin in mediating the binding of algae to the progenitor endosymbiotic cells to aid the interaction with algae in the stalk-mouth region of the gastrodermis.

### Identification of scavenger receptors and actin regulators for endosymbiosis

Upon binding to the endosymbiotic algae, coral cells must activate the phagocytic pathway for alga uptake. As a secreted protein, LePin needs to use phagocytic receptors to initiate phagocytosis. We examined genes in the GEP1, 4, and 22 and found a number of genes predicted to encode scavenger receptors (Supplementary Table S2). The scavenger receptors are known to recognize common ligands on microbial surfaces and are implicated in initiating microbial phagocytosis^32,33^. We performed shRNA treatment and found that shRNA targeting of each of two scavenger receptor genes, *CD36* and *DMBT1*, resulted in a significant reduction of the newly formed alga-containing endosymbiotic cells (Fig. 5a, Extended Data Fig. 2c). Both *CD36* and *DMBT1* were previously identified as marker genes for the *Xenia* endosymbiotic cell type^7^. Additionally, sequence analyses showed that the *Xenia* CD36 has an N- and C-terminal transmembrane domains similar to the CD36 known to initiate microbe phagocytosis and host immune response in the well-established model organisms^34,35^. DMBT1 has an established role in modulating immune response in the mucosal surfaces in mammals^36,37^. Thus, these two scavenger receptors may function downstream of LePin to engage the phagocytic machinery and modulate *Xenia* immunity for algae uptake.

**Fig. 5.**
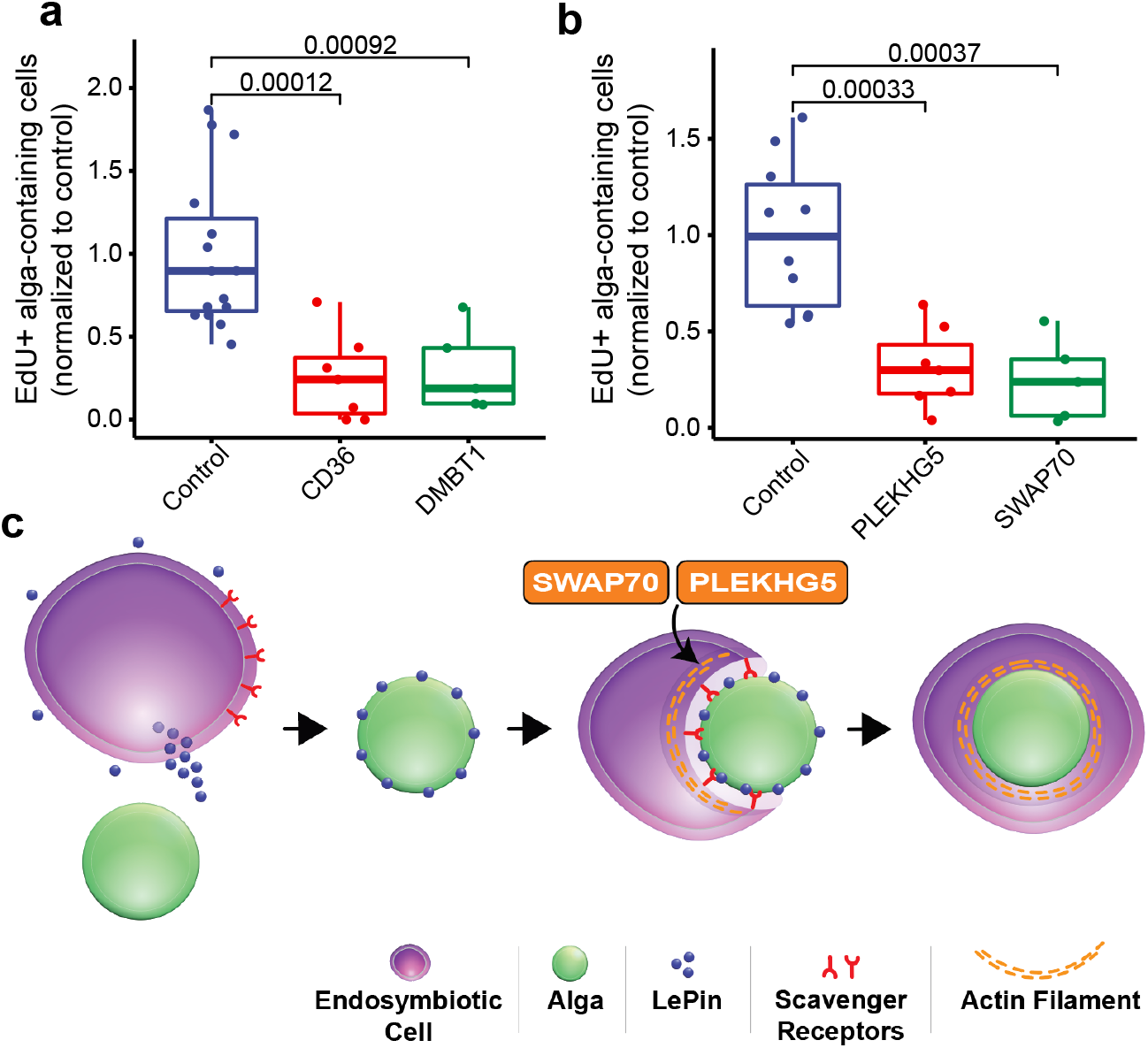
scavenger receptors and actin regulators for endosymbiosis. **a, b**, shRNAs targeting scavenger receptors, CD36 and DMBT1 (**a**), or actin regulators, PLEKHG5 and SWAP70 (**b**), reduced the number of newly formed alga-containing endosymbiotic cells. Quantification is the same as in Fig. 2**d**. Each dot stands for a polyp. 5-9 polyps were used for each shRNA treatment. The p values of the unpaired Student’s t-test are indicated. **c**, A model: the secreted LePin expressed in the endosymbiotic cells binds to alga and initiates alga recognition. This leads to the engagement of the scavenger receptors CD36 and DMBT1 and alga phagocytosis mediated by the actin regulators PLEKHG5 and SWAP70.

To establish endosymbiosis, *Xenia* needs to not only phagocytose the endosymbiotic algae, but also repress the fusion of the alga-containing phagosomes with lysosomes, thereby supporting the conversion of phagosomes to symbiosomes^38^. Actomyosin assembly is required for phagocytic target search and capture, formation of phagocytic cup, the subsequent phagosome formation and maturation^39^. Additionally, the disassembly of actin cables on the phagosome surfaces can aid phagosome maturation through fusion with endosomes and lysosome^40^. By surveying the GEP1, 4, and 22, we found two genes encoding conserved actin regulators, PLEKHG5 and SWAP70, in GEP4 and 22 (Supplementary Table S2, Extended Data Fig. 2d). PLEKHG5 is a marker for endosymbiotic cells and as a RhoGTPase guanine nucleotide exchange factor (RhoGEF), PLEKHG5 can regulate actomyosin assembly at the plasma membrane^41^. SWAP70 stimulates actin cable formation on phagosome membranes during phagocytosis and may inhibit phagosome maturation and fusion with lysosomes^42^. We found that RNAi of either gene resulted in a significant reduction of newly formed alga-containing endosymbiotic cells (Fig. 5b). This suggests that the two proteins are part of the phagocytic machinery functioning downstream of scavenger receptors to enable algae phagocytosis. By favoring actin cable assembly on the phagosome surface^42^, SWAP70 could delay phagolysosome fusion and aid symbiosome formation in *Xenia*.

### Summary and outlook

Through developing an efficient RNAi protocol, we have identified a secreted lectin, LePin, that supports the formation of alga-containing endosymbiotic cells in *Xenia*. Since LePin interacts with algae and appears to facilitate their binding to the stalk and mouth region of *Xenia* gastrodermis, it may function in the selection of specific *Symbiodiniaceae* species for *Xenia* endosymbiosis. It is known that corals use a select set of *Symbiodiniaceae* species for endosymbiosis and the corals that select the heat resistant *Symbiodiniaceae* exhibit resistance to heat-induced bleaching^15,43^. Since lectins can recognize specific glycans found on algal surface, further comparative studies of LePin-like lectins in various corals and their cognate *Symbiodiniaceae* may lead to identification of the LePin sequence motif that selects for heat resistant algae. With the availability of CRISPR (Clustered Regularly Interspaced Short Palindromic Repeats) for coral gene editing^44^, it could then be possible to engineer heat sensitive corals by editing LePin to allow their selection for heat resistant algae. This could offer a possible biological mitigation for coral reefs in the face of global warming.

We also demonstrate the power of applying bioinformatics tools to identify gene expression programs that contain important regulators of endosymbiosis. These genes could serve as a great resource for the field for discovery of additional regulators of endosymbiosis. By focusing on the GEPs whose expressions are influenced by LePin during early and mid-stages of endosymbiosis, we identified genes that function during early steps of alga uptake (Fig. 5c). With two transmembrane domains, the *Xenia* CD36 we report here is likely to trigger phagocytosis of the LePin bound algae attached to the newly divided endosymbiotic cells. CD36 may also regulate *Xenia* immune response via its N and C-terminal cytoplasmic sequences because the CD36 in the well-established model organisms is known to engage various innate immune pathways upon microbe binding and phagocytosis. *Xenia* and mammalian DMBT1 are highly similar in their multiple scavenger receptor domains. Mammalian DMBT1 can suppress inflammation^36^. Further study of these *Xenia* scavenger receptors could provide mechanistic insights on algae uptake and immune modulation critical for endosymbiosis. The *Xenia* PLEKHG5 and SWAP70 are conserved actin regulators that can mediate phagocytosing algae and phagosome formation. Since symbiosomes are derived from phagosomes and since SWAP70 is implicated in delaying phagosome maturation in mammalian cells, further study of the *Xenia* SWAP70 could facilitate the understanding of how symbiosomes are derived from phagosomes to aid nutrients exchange between coral and algae.

---

**Extended Data Fig. 1.**
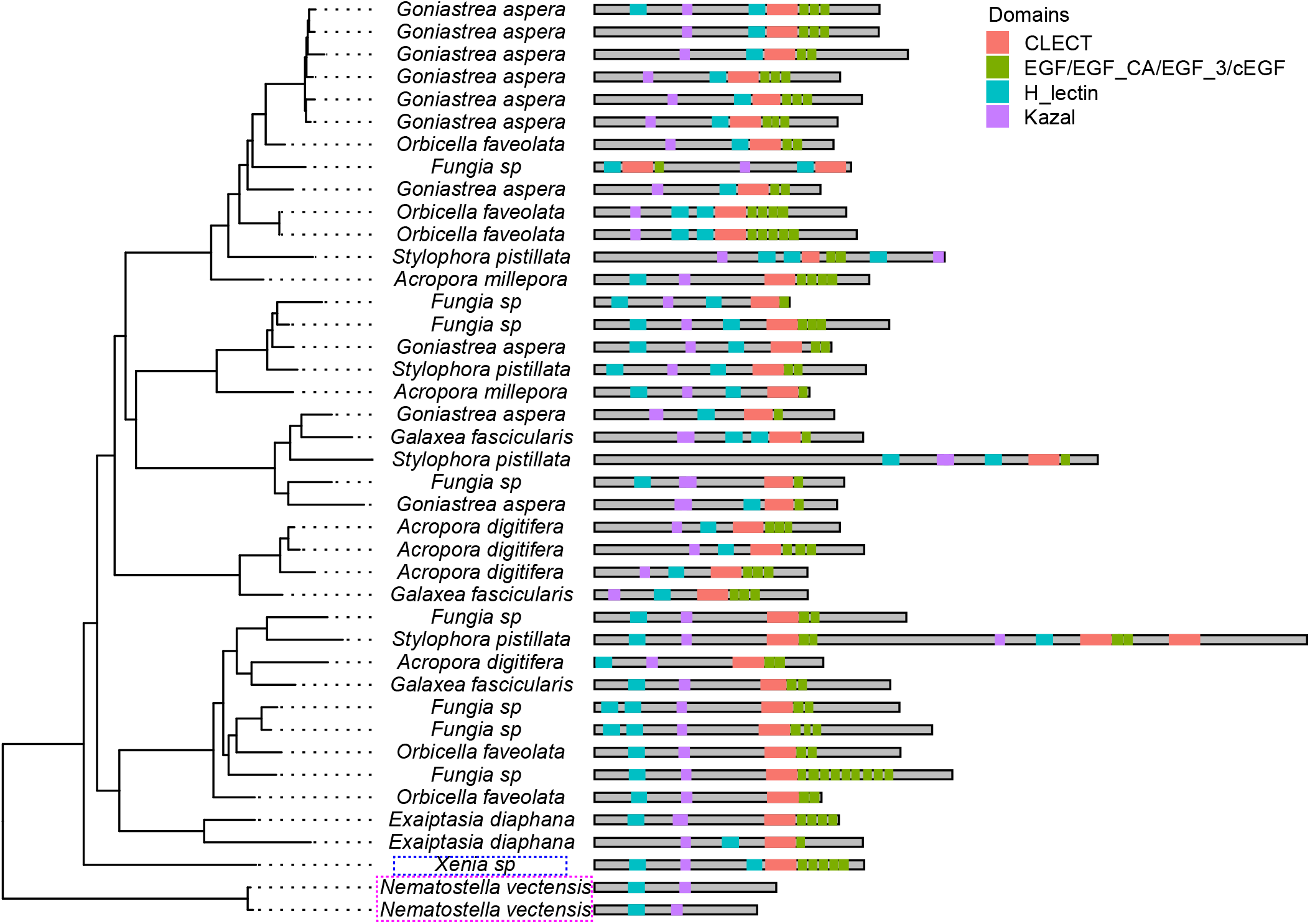
Domain organization of the predicted LePin-like proteins from different marine cnidarians plotted with the phylogenetic tree based on LePin sequences. CLECT: C type Lectin domain; EGF/EGF_Ca/EGF_3/cEGF: EGF, EGF-Cacium binding and EGF like domains (EGF_3 or cEGF); H_lectin: H type lectin domain; Kazal: kazal domain. *Xenia* LePin is highlighted by a dashed blue rectangle. The two lectins in Nematostella vectensis (highlighted by a dashed magenta rectangle) that are most similar to *Xenia* LePin miss a few domains and appear as an outgroup.

**Extended Data Fig. 2.**
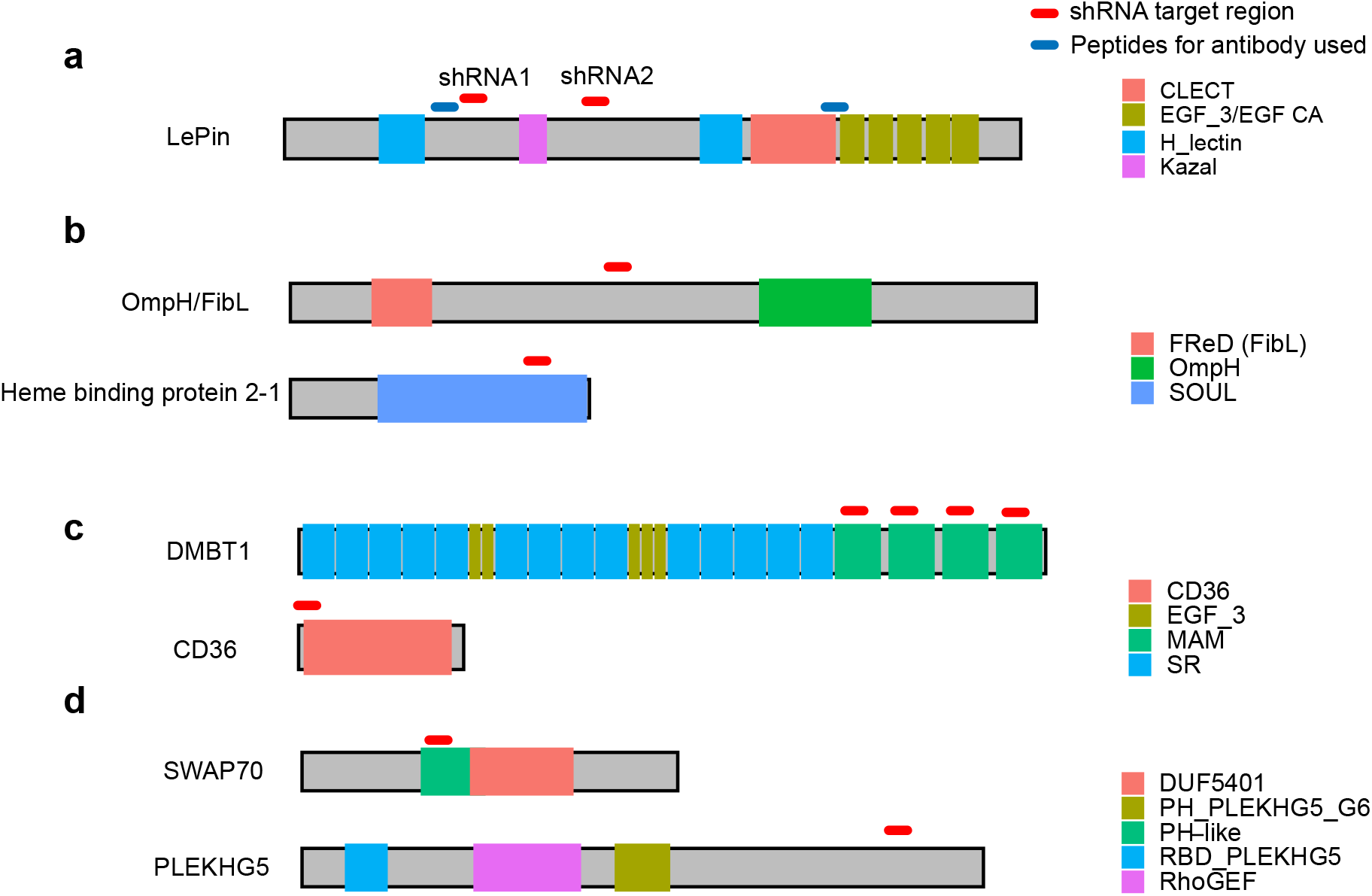
Illustration for proteins studied. Illustrations of domains and target regions of shRNA and peptides (for antibodies) for the proteins studied in this report.

**Extended Data Fig. 3.**
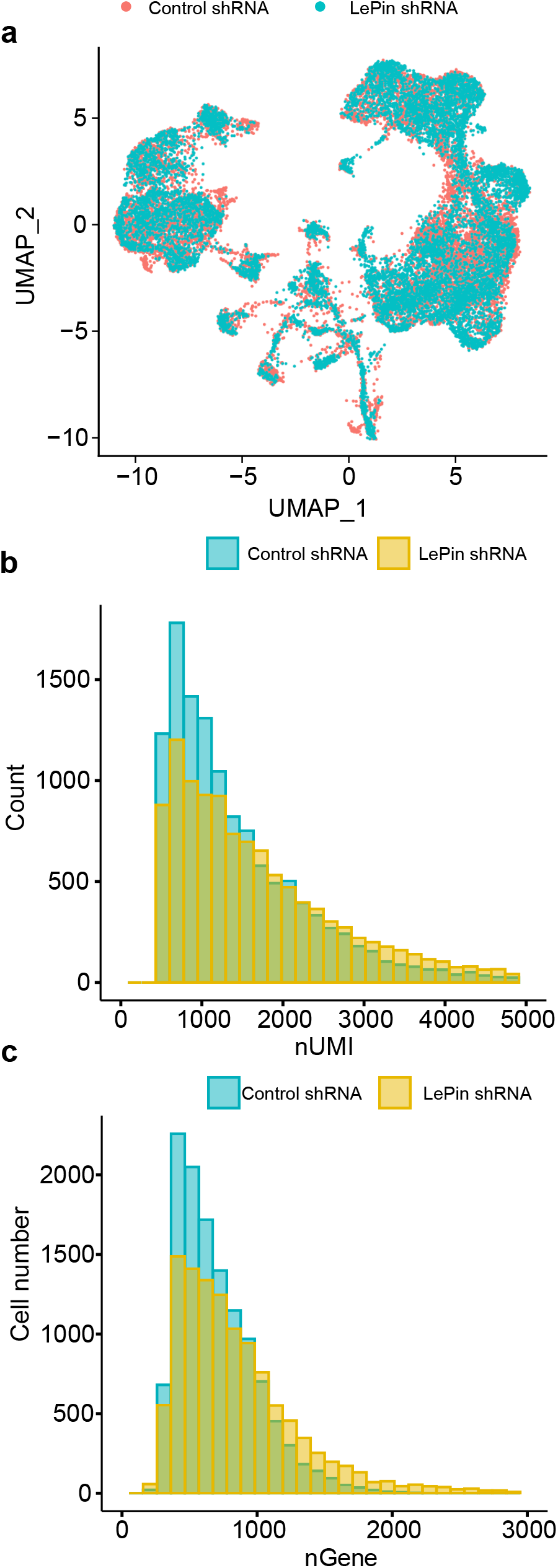
Additional analyses of scRNA-seq for RNAi-treated samples. **a,** The integrated UMAP of all cells from the control and LePin RNAi samples. **b, c**, Distributions of the detected UMI (Unique Molecular Identifier) numbers (**b**) and gene numbers (**c**).

**Extended Data Fig. 4.**
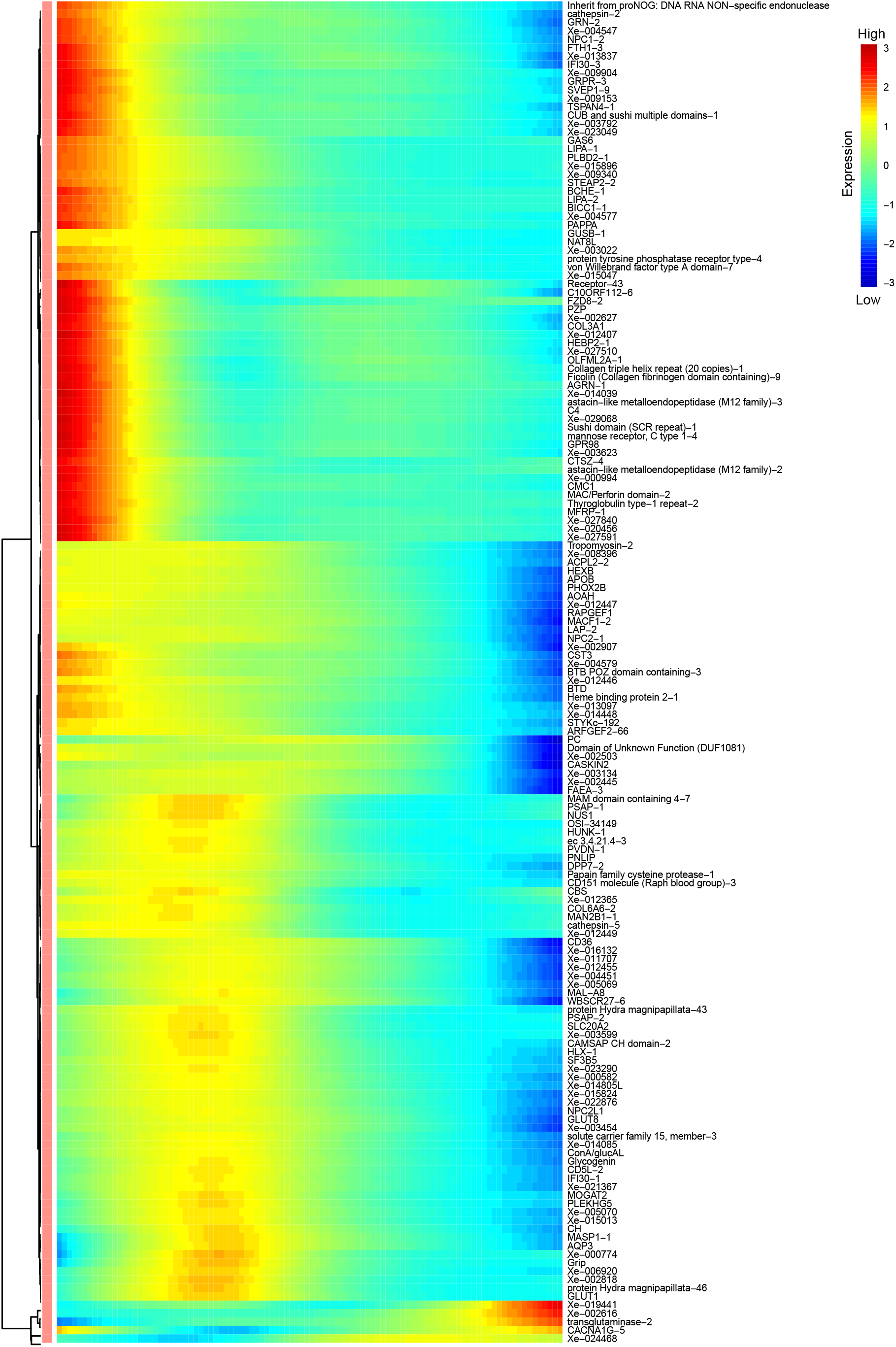
An expanded view of Fig. 2b to show the names of individual genes.

**Extended Data Fig. 5.**
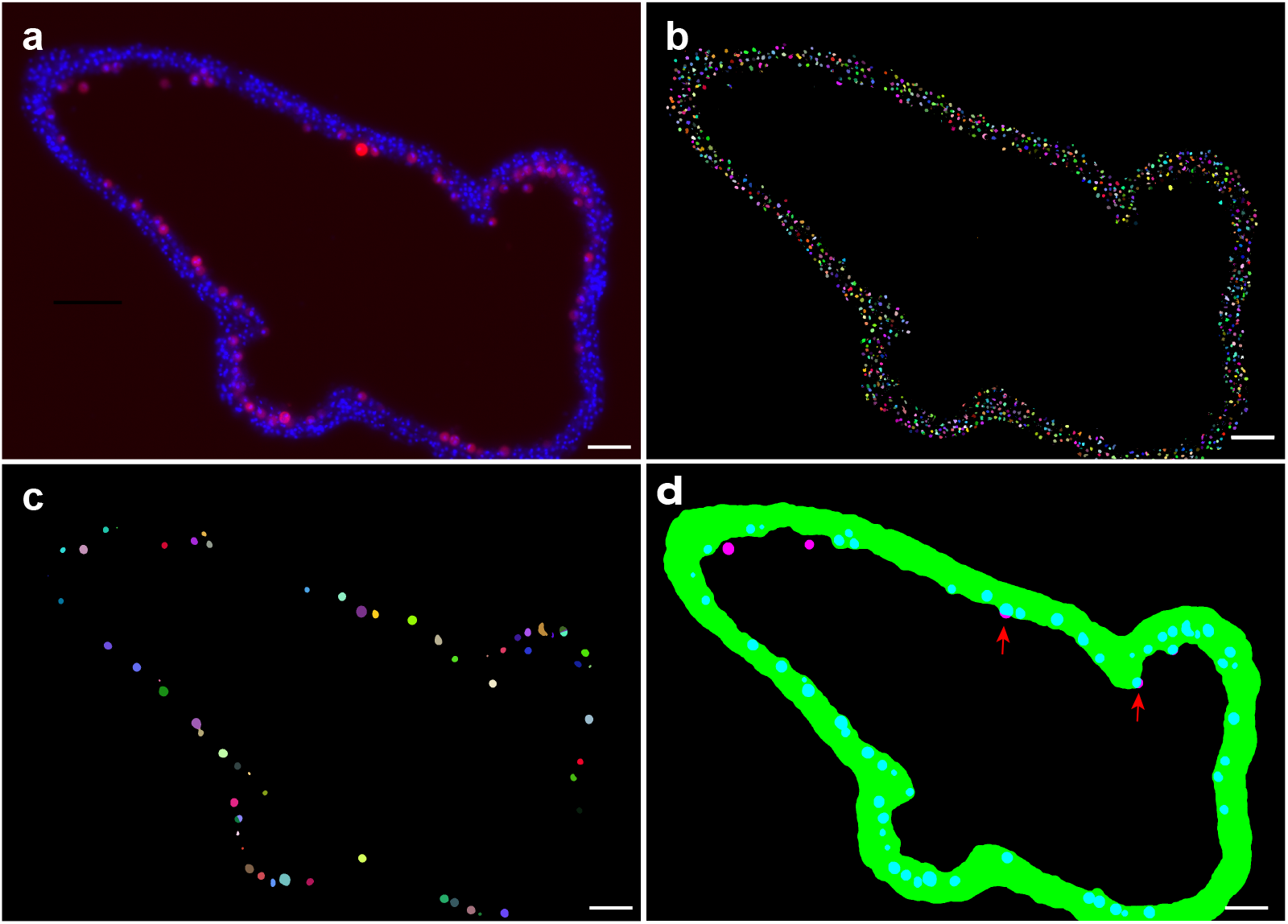
Unbiased high throughput quantification of algae attached to the surface of *Xenia* gastrodermis. **a,** An example of an epifluorescence image from a tissue section. Blue, DAPI staining of nuclei. Red, auto-fluorescence from the alga. **b**, Labeling of all cells by the DAPI signal. Each nucleus was pseudo colored to enable counting (see the Method section). **c**, Labeling of alga cells by the auto-fluorescence signal. Each alga was pseudo colored to enable counting (see the Method section). **d**, Tissue mask (green) is generated with a lower threshold of the DAPI signal and overlaid with the algal autofluorescence channel. The algae within the tissue mask are labeled blue. The algae outside or not completely inside the tissue mask are labeled pink. Red arrows point to the algae protruding from the tissue surface and are counted as algae attached to the surface of the gastrodermis. Scale bar, 50μm

**Extended Data Fig. 6.**
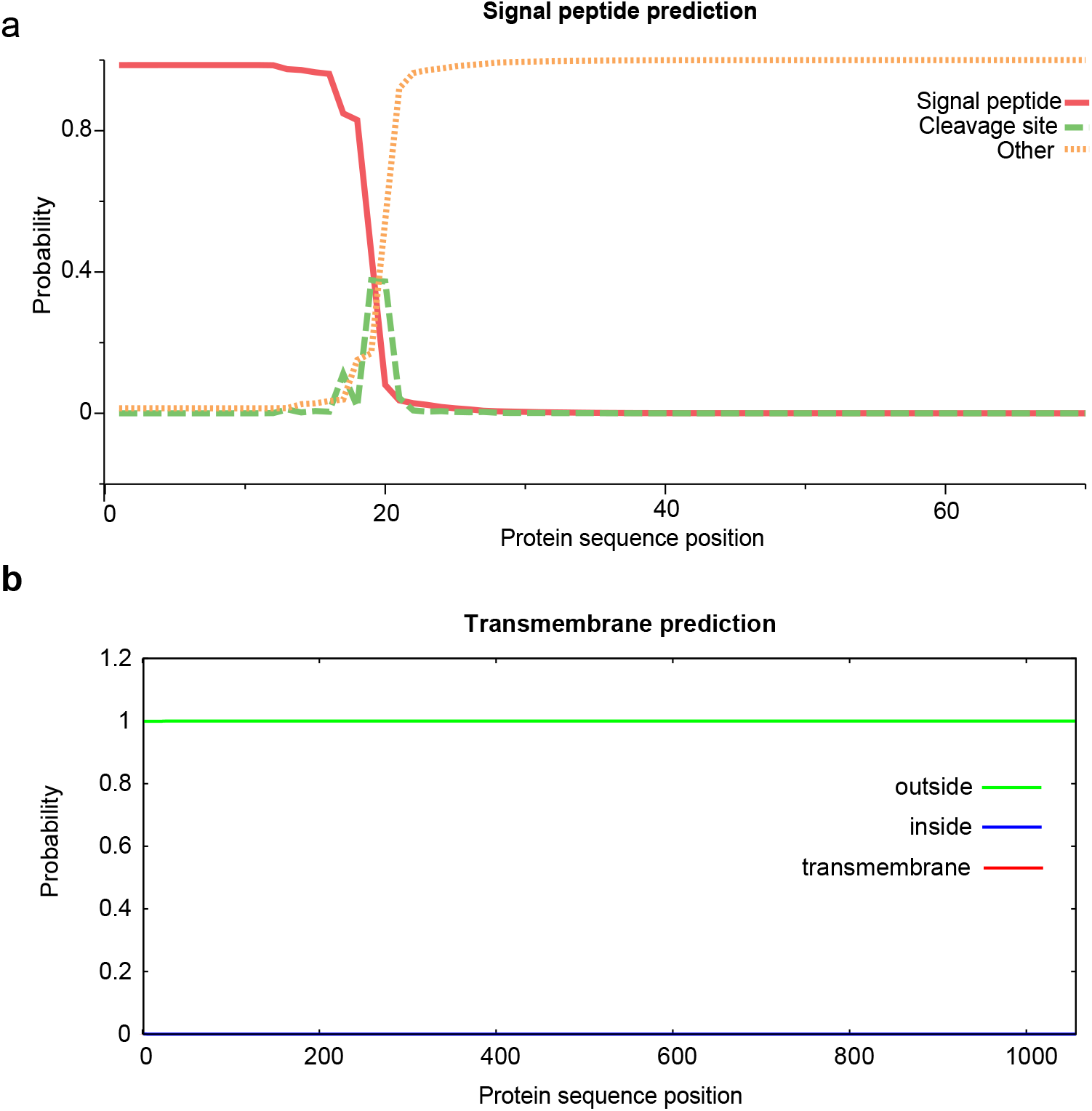
Prediction of signal peptide and transmembrane sequences in LePin. **a,** the signal peptide is predicted by SignalP 5.0 with the Eukarya model. OTHER: no signal peptide predicted. Only the first 70 amino acids of LePin are plotted. **b**, LePin is predicted as a protein without transmembrane domain. TMHMM (v2.0) was used for the prediction. The full length LePin sequence is plotted.

**Extended Data Fig. 7.**
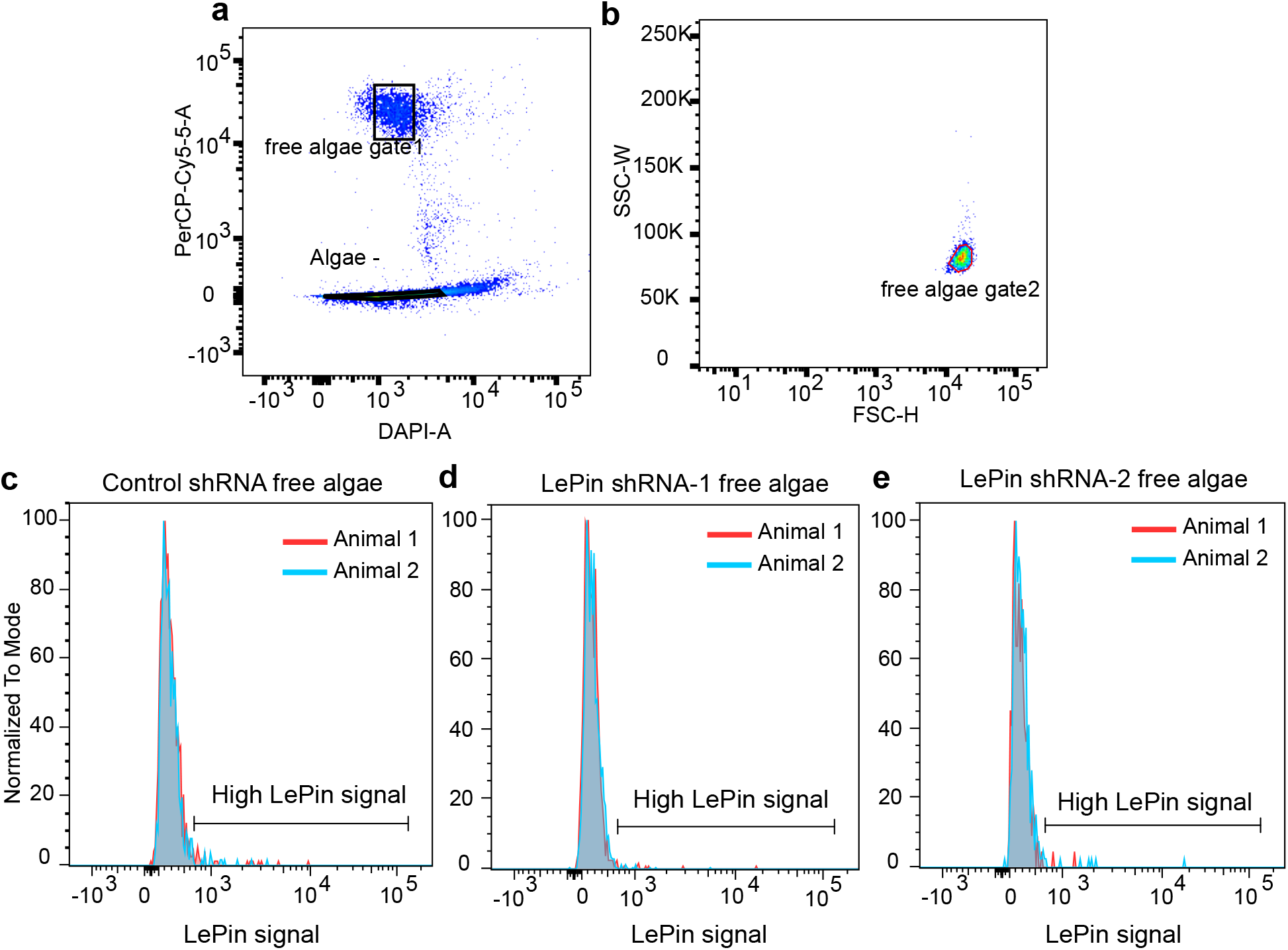
Quantification of LePin signal on the isolated free algae in *Xenia* by FACS. **a,b,** Gating strategy for the free algae in *Xenia*. The algae are first gated based on the DAPI staining of the nuclei and algae autofluoresce (Cy5.5 signal) (**a**, free algae gate1). The algae are further gated based on the forward scatter (FSC) and side scatter (SSC) signals to further exclude those algae that are inside the *Xenia* cells (**b**, free algae gate2). **c-e**, LePin signal distribution on free algae in control (**c**) and LePin knocking down samples (**d,e**). Color codes for individual animals. The percentage of algae with high LePin signals (as indicated by the brackets) are quantified.

## Methods

### *Xenia* culture and regeneration

The *Xenia* colonies are grown in our aquarium as previously described^45^. To culture polyps in tissue culture plates, one or several colonies of *Xenia* is collected from the aquarium and incubated in the Ca-free seawater (393.1 mM NaCl, 10.2 mM KCl, 15.7 mM MgSO_4_·7H_2_O, 51.4 mM MgCl_2_·6H_2_O, 21.1 mM Na_2_SO_4_, and 3 mM NaHCO_3_, pH 8.5) for 15 min to relax the animals. Individual polyps are cut from the colony and transferred into a well of 24 well plates with 2 ml of seawater. Colony health is important for good regeneration. High polyp lethality rates after transplanting to tissue culture plates generally lead to poor regeneration. We only perform the follow up experiments and analyses if the survival rate is >80%. The plates are incubated at 25°C in a Percival incubator with a cycle of 12 hours light on and 12 hours of light off. The polyps are allowed to recover for a week after transplantation before performing the follow up experiments. For *Xenia* regeneration, all 8 tentacles are surgically cut away and the stalk attached to the plate is allowed to regenerate by further incubation.

### shRNA design and production

The shRNA target sequences are designed using the Invitrogen’s siRNA Wizard site (https://www.invivogen.com/sirnawizard/design.php) with 19 nucleotides as the motif length, targeting a given gene. The 19 nucleotide sequences selected have 40-50% GC content and no more than four consecutive thymidines (T). These selected candidate sequences are then blasted against the *Xenia* transcriptome to select the ones with specific hits to the target gene. The T3 promoter sequence (5′AATTAACCCTCACTAAAG 3′) is added to the 5’ of these targeting sequences followed by adding the loop sequence (5’-TTCAAGAGA-3’) to the 3’ end of the target sequence and then the reverse complementary sequence of the target sequence. Two additional Ts are then added to the 3’ end of the synthetic sequence, which gives rise to the shRNA primer1. Both shRNA primer1 and its reverse complementary strand (shRNA primer2) are synthesized as DNA oligos. To confirm the stem loop structures, we used the mfold Web Server (http://unafold.rna.albany.edu/?q=mfold/RNA-Folding-Form) to evaluate stem-loop structures. The two complementary DNA oligos are mixed in equal molar (0.2 pM each for 20μl reaction) and used as the template for *in vitro* transcription by MEGAscript™ T3 Transcription Kit (Invitrogen, AM1338). After transcription, 1μl DNase I (Ambion™) is added for each 20 μl reaction. The DNA templates are removed by incubating at 37 °C for 30 min. To purify the shRNA, one-tenth volume of 3 M NaOAc (pH 5.5), and 3 volumes of ice-cold 95% ethanol are added into the reaction. shRNA is precipitated at −80°C for 30 min, followed by centrifuging at 14,000 rpm for 15 min at 4°C. The pellet is washed twice with 500μl ice-cold 80% ethanol and air dried at room temperature. The shRNA is further dissolved in DEPC-treated water and the concentration is determined by Nanodrop. The sequence information for all the shRNA oligos used in this study is listed in Supplementary table 1 and the target positions of these oligos in the corresponding proteins are indicated in Fig. S2.

### shRNA treatment of *Xenia* polyps

Each intact polyp or regenerating polyp is cultured in a well of 24 well plates. After tentacle amputation, healthy polyps under good culturing conditions grow a clear set of short tentacles by day 3 and the tentacles continue to extend in subsequent days. The polyps with regeneration delays are excluded for RNAi experiment. For RNAi treatment, the sea water in each well is replaced with 500 μl fresh sea water containing 20 μg shRNA. The animals are incubated for 3 hours followed by adding an additional 500 μl fresh sea water. The animals are further incubated for 3 days before further analyses. When healthy intact polyps or regenerating polyps are used, we have not observed death caused by shRNA treatment.

### Edu pulse-chase and shRNA treatment of regenerating *Xenia* polyps to analyze the newly formed alga-containing endosymbiotic cells

The tentacle amputated *Xenia* polyps are allowed to regenerate and 1 mM EdU is added to label the proliferating cells on day 3 and incubated for 48 hours. Edu is washed away using sea water and the polyps are incubated further. The cells that contain algae and are EdU positive are newly formed alga-containing endosymbiotic cells. These cells are quantified by FACS as previously described^1^. Our previous studies have shown that the number of newly formed alga-containing endosymbiotic cells continue to increase until the regeneration day 15^45^. The regenerating polyps are then treated with shRNA on day 6. FACS analysis is performed four days after shRNA addition at the end of regeneration day 9.

### Single cell RNA sequencing (scRNA-seq)

The scRNA-seq is carried out using the 10x Genomics platform as we previously described^1^. Briefly, two regenerating *Xenia* polyps treated by control RNAi or LePin RNAi from regeneration day 6 to the end of regeneration day 9 (4 days) are digested with 1 ml digestion buffer, containing 3.6 mg/ml dispase II (Sigma, D4693), 0.25 mg/ml liberase (Sigma, 5401119001), 4% L-cysteine in Ca-free seawater. After one hour incubation at room temperature, the polyps are dissociated into single cell suspension. After tissue dissociation, fetal bovine serum is added to a final concentration of 8% to stop enzymatic activities. The cell suspension is filtered through a 40-μm cell strainer (FALCON) twice. Approximately 17,000 cells per sample are used for single-cell library preparation using the 10× Genomics platform with Chromium Next GEM Single Cell 3′ GEM, Library and Gel Bead Kit v.3.1 (PN-1000121, v.3 chemistry), Chromium Next GEM Chip G Single Cell Kit (PN-1000127), and i7 Multiplex Kit (PN-120262) according to manufacturer’s manual. Libraries are sequenced using Illumina NextSeq 500 for paired-end reads. In order to reduce batch effect, the control and LePin RNAi treated samples are loaded in two wells in one 8-well strip and sequenced at the same time. Sample preparations are carried out simultaneously with multiple channel pipette.

### LePin antibody generation, purification, and antibody immunofluorescence microscopy

Three LePin peptides, KHNTSNNNVDPKYNA, NKNHPTSENNYAYCK, KNYTCQKDYDECGEN, are selected based on the high antigenicity and surface probability of the LePin protein sequence. A cysteine is added to the N terminus of each peptide for coupling chemistry and the peptides are synthesized by GenScript (Inc.). The purity of the three peptides are 96.9%, 90.5% and 85.3%, respectively. The antibodies are produced at Pocono Rabbit Farm & Laboratory (Inc.) using the 28 day mighty quick protocol. The second and third peptides are combined and injected into two rabbits (#36232, #36243), while two additional rabbits (#36230,#36231) are immunized by the first peptide. To assess the sera, each is used for immunostaining of *Xenia* tissue sections. The sera from three rabbits (#36231, #36232, #36243) exhibit stronger immunofluorescence signals than the corresponding control pre-immunization sera.

The LePin antibodies from the serum of rabbit (#36232) is affinity purified using the corresponding LePin peptides (NKNHPTSENNYAYCK and KNYTCQKDYDECGEN). Briefly, the LePin peptides are coupled to an affinity column with SulfoLink® Immobilization Kit (Thermo Scientific™, Cat. 44995) according to the manufacturer’s manual. The serum is slowly loaded onto the column twice over at least 2 hours. The column is washed with 10 column volumes of Tris-buffered saline (TBS, 20 mM Tris, 150 mM NaCl, PH7.4), followed by additional washing with 10 column volumes of the washing buffer containing 0.5 M NaCl, 20 mM Tris HCl, pH 7.4, 0.2% Triton X-100. After one more washing with 10 column volumes of washing buffer, the antibody is eluted with 0.2 M glycine, pH 2.0. The eluate is collected at 3 ml/fraction into tubes containing 0.2 ml 2 M Tris base (pH 11). The protein concentration of each fraction is measured at OD280. The fractions with OD280>0.1 are pooled and dialyzed against 2L TBS overnight at 4°C. The antibody solution is further concentrated by placing the dialysis bags in Aquacide. Aliquots of 0.5 ml of the concentrated antibody are snap frozen in liquid nitrogen and stored at −80°C. Upon usage, an aliquot is thawed at room temperature and further concentrated with a 100 kDa Ultra-0.5 Centrifugal Filter Unit (Amicon^™^) to a concentration of 2.2μg/μl.

The antibody staining is carried out according to the previous publication^46^ with minor modifications. Briefly, the polyps are pretreated with 2% ice-cold HCl for 3 minutes to remove mucus^47^ and then fixed with 4% paraformaldehyde (PFA) for 15 min at room temperature. After washing twice with PBST (0.5% triton X-100 in PBS: 137 mM NaCl, 2.7 mM KCl, 10 mM Na_2_HPO_4_, and 1.8 mM KH_2_PO_4_), the polyps are transferred to 30% sucrose and incubated overnight at 4°C. Polyps are then embedded in Tissue-Tek O.C.T. and subjected to cryosection at 12 μm thickness. The slides containing tissue sections are rehydrated by incubating with PBST for 10 min for three times. The slides are then blocked with 10% goat serum (Sigma, G9023) in PBST for 40 min at room temperature. The affinity purified LePin antibodies from rabbit #3264 or rabbit IgG isotype control (invitrogen, Cat. 10500c) are diluted to 22 ng/μl with 10% goat serum and used to incubate with the slides overnight at 4°C. After washing with PBST three times 10 min each, the secondary antibody (Goat anti-Rabbit IgG, Invitrogen, 1:400 dilution,) and DAPI are applied to the slides. The slides are incubated at room temperature for 1 hour, followed by washing with PBST three times, 10 min each. Slides are mounted with ProLong™ Gold Antifade Mountant (Invitrogen™, P36930). The images are acquired with a confocal microscope (Leica).

### LePin phylogenetic analyses

To build LePin phylogenetic tree among marine anthozoans with available sequence information, Blastp(v2.1.2) is used to find the potential *LePin*-like genes with parameter -max_target_seqs 5000 -max_hsps -evalue 1e-10. The homologous regions in a given protein sequence that cover less than 60% of the *Xenia* LePin are filtered out. For the marine anthozoans known to do endosymbiosis with algae, the genes without the predicted KAZAL, EGF, CLECT, H_lectin domains are also excluded. The phylogenetic tree is built by ETE toolkit (v.3) with the standard_trimmed_fasttree workflow^48^. Domain information for each gene is fetched from NCBI CDD domain database (April 02, 2019 version)^49^.

### NMF analyses of the scRNA-seq datasets

scRNA-seq analysis is carried out as previously described with minor modifications^45^. Briefly, the gene expression matrix for each cell is generated by cellranger(v3.1). Cells with mitochondrial gene expression higher than 0.2% of all reads or UMI values lower than 400 were excluded. Potential doublets as identified by DoubletFinder (v2.0.3)^50^ are excluded before subsequent analysis. The endosymbiotic cells from LePin and control RNAi samples are identified by Seura(v3.0.2)^51^ label transfer method using our previous *Xenia* scRNA-seq data^45^ as reference and subject to monocle2 (v2.12.0)^52^ for lineage trajectory prediction.

The NMF analysis is carried out according to the previous publication^53^. Briefly, we applied the NMF analysis on the scRNA-seq data for the endosymbiotic cell lineage. To select the highly variable expression genes, the 3000 variable genes from control or LePin RNAi are identified with Seurat FindVariableFeatures functions. These genes are combined together which leads to a total of 5194 variable genes. The NMF analysis is based on previously published NMF framework (https://github.com/YiqunW/NMF) and performed with log-normalized expression of the variable genes in all endosymbiotic cells using the run_nmf.py function with the default parameters. K=45 is selected and the Gene Expression Programs (GEP) with low reproducibility between different runs excluded as previously described^53^. The NMF analysis will generate two matrices. One is GEP-gene matrix, which shows the relative contribution of each gene (also called gene weight) to the GEP. The other is the GEP-cell matrix, which shows the relative expression of GEP (also called GEP weight) to each cell. The GEPs that only contain genes with low weight (<0.6) are also excluded from further analysis of the effect of LePin in GEP expression along the endosymbiotic lineage development. The relative expression of each GEP is used to indicate the GEP activity for each cell and plotted along the pseudotime trajectory (Supplementary Table S3).Cells are grouped into control or LePin RNAi samples to further explore the GEP activity change upon LePin RNAi.

## Data availability

We have uploaded all raw data to NCBI (PRJNA869069). The *Xenia* genome is also available at NCBI (Genebank accession: JAJSDR000000000.1). Select intermediate RDS objects are available at figshare (https://figshare.com/articles/dataset/Processed_R_objects_for_LePin_RNAi_/20481900).

## Code availability

All analysis codes for scRNA-seq and alga counting are available at GitHub: https://github.com/MinjieHu/Xenia_RNAi.

## Acknowledgements

We thank Fred Tan and Alison Pinder for assistance with all the sequencing and initial processing of raw reads; Navid Marvi for the model sketch; and Lynne Hugendubler and Mike Watts for maintaining the coral aquarium; Ross Pedersen and Joseph Tran for critical comments. This work is supported by Gordon and Betty Moore Foundation, Aquatic Symbiosis no. GBMF9198 (https://doi.org/10.37807/GBMF9198, Y.Z.).

## Author information

Authors and Affiliations

Department of Embryology, Carnegie Institution for Science, Baltimore, MD, USA

Minjie Hu, Yun Bai, Xiaobin Zheng & Yixian Zheng

## Contributions

M.H.and Y.Z. conceived the project. M.H. and Y.Z. designed experiments. M.H. and Y.B. performed the experiments. M.H. and X.Z. analysed the data. M.H., Y.B., X.Z. and Y.Z. interpreted the data. M.H. and Y.Z. wrote the manuscript.

## Additional information

Supplementary Information is available for this paper.

---

**Supplementary Table S1. The list of genes/proteins studied**. The shRNA and predicted protein sequences are listed along with the gene names. We manually checked each gene to confirm that it has reads mapping to all their exons using transcriptomic data.

**Supplementary Table S2. The top 30 genes for each of the 24 GEPs.** The first sheet contains the list of 30 genes in each GEP. Each GEP has two columns. The first column shows the top 30 genes. The second column indicates the weight of each gene. The second sheet (related gene id and sequence) shows the protein sequence and the gene ID of the genes shown in the first sheet.

**Supplementary Table S3. Expression of each GEP along the pseudotime for each endosymbiotic cell.** The cell identity comes from the cell barcode together with sample information. The pseudotime value for each cell is calculated by the lineage trajectory analysis with Monocle2. The relative expression of each GEPs is calculated by NMF analysis.

